# Unsupervised temporal consistency improvement for microscopy video segmentation with Siamese networks

**DOI:** 10.1101/2021.03.25.436993

**Authors:** Akhmedkhan Shabanov, Daja Schichler, Constantin Pape, Sara Cuylen-Haering, Anna Kreshuk

## Abstract

We introduce a simple mechanism by which a CNN trained to perform semantic segmentation of individual images can be re-trained - with no additional annotations - to improve its performance for segmentation of videos. We put the segmentation CNN in a Siamese setup with shared weights and train both for segmentation accuracy on annotated images and for segmentation similarity on unlabelled consecutive video frames. Our main application is live microscopy imaging of membrane-less organelles where the fluorescent groundtruth for virtual staining can only be acquired for individual frames. The method is directly applicable to other microscopy modalities, as we demonstrate by experiments on the Cell Segmentation Benchmark. Our code is available at https://github.com/kreshuklab/learning-temporal-consistency.

## 1. INTRODUCTION

Live imaging is a cornerstone microscopy technique of molecular biology, with applications ranging from analysis of sub-cellular processes to morphodynamics of tissues and organisms. Automatic processing of live imaging videos usually includes segmentation and tracking tasks, which for many cutting-edge biology problems turn out to be quite challenging due to low contrast and signal-to-noise ratios as low fluorophore excitation level is necessary to avoid photo-bleaching and phototoxicity. Furthermore, for many delicate processes the addition of fluorescent proteins is known to change the cell phenotype. The imaging thus needs to be performed label-free, relying on the difference of compartment refractive index to produce contrast. Recently, several methods have been proposed to address this problem by the so called ”virtual staining”: training of a convolutional neural network (CNN) to either generate a staining from label-free images or to segment label-free images based on groundtruth images with staining [1, 2, 3]. These methods turned out to be so successful that the ability to generate labeled groundtruth is now built into several commercial microscopes.

However, some cellular processes are so sensitive to light that for those the fluorescent labeled groundtruth cannot be acquired as video. Common examples include the studies of membrane-less organelles - dynamic phase-separated structures that are maintained through multiple weak and multivalent interactions [4]. To avoid alteration of phase separation dynamics by light or the fluorescent tag [5], the training data needs to be acquired as still frames and the segmentation algorithms - which will later be applied to live imaging recordings - cannot directly exploit temporal context. A similar concern arises for the general microscopy video segmentation task in the frequent setting where little manually labeled groundtruth is available. For example, thanks to the Data Science Bowl Kaggle challenge [6] ample groundtruth is now available for the 2D nuclei segmentation task. However, a network trained on this groundtruth has no notion of temporal consistency and cannot in any way exploit the temporal nature of the data when it is applied to videos.

The aim of our contribution is to close this gap by an unsupervised extension to the training of the segmentation network. A Siamese network is used to connect video segmentation trained to maximize temporal consistency between unlabeled frames and regular image segmentation trained on labeled snapshots (Fig. 1). We demonstrate how our setup can be used to improve the segmentation of nuclei and nucleoli in long or fast time-lapse holotomographic movies where parallel fluorescent imaging is not possible due to photobleaching or is avoided as to not alter the properties of the cellular structures. Building on this result, we demonstrate how our approach can be used to improve a general-purpose segmentation network for prediction on videos. To that end, we retrain the popular StarDist method [7] with annotations from the Data Science Bowl to segment fluorescent videos from the Cell Tracking Challenge [8], obtaining a significant improvement in average precision score.

**Fig. 1.**
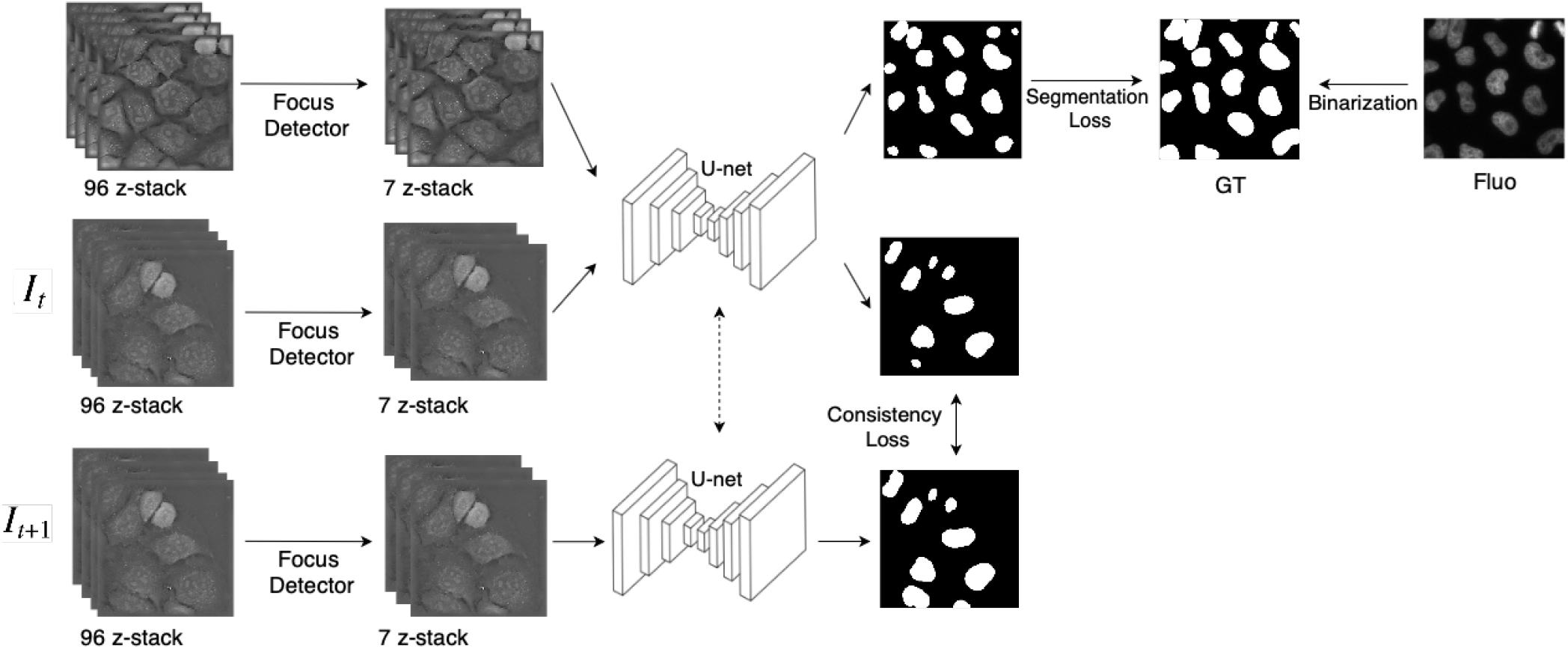
Architecture of our setup as applied to the holotomographic imaging task with training labels from parallel imaging with fluorescent markers for individual frames.

## 2. METHOD

We identify our task as segmentation of completely unlabeled videos using independent individual labeled images as training data. While multiple methods have recently been introduced for segmenting videos with a few or just one labeled frame [9, 10, 11, 12], none of them are directly applicable to our use case. We have therefore developed a new approach, the core of which lies in training on both labeled and unlabeled data through the Siamese duplication of the segmentation network.

Siamese networks have originally been introduced for the object tracking task, where the object bounding box was identified in the first frame of the video [13]. The architecture allowed to train the network offline to predict similarity between image patches. This approach has later been extended to simultaneous object segmentation and tracking[14] by adding another branch that was trained to segment, while the similarity loss was applied to patches as before. The network was trained on millions of annotated video frames and implicitly learned temporal dynamics and expected temporal consistency in natural videos. No annotated datasets of such size exist for live microscopy imaging, but as the imaging experiments are targeted and highly controlled, prior biological knowledge can be exploited to estimate and then impose temporal consistency rules. In our approach, this step is realized through an additional loss on the similarity of the segmentations of consecutive frames of unlabeled videos. We assume that the captured biological processes have a certain degree of temporal consistency and segmentations of consecutive time frames are are not completely dissimilar.

Our full setup is shown in Fig. 1. As the backbone, we take the standard U-net [15] trained to segment 2D images and make a Siamese network of 2 U-nets with shared weights.

The training proceeds as follows in alternating steps:

1. Draw a batch of annotated 2D images, predict their segmentation.
2. Compute the DICE loss [16] (or any other semantic segmentation loss) wrt the labels and back-propagate through one of the Siamese sub-networks
3. Draw a batch of unlabeled consecutive video frame pairs, predict their segmentations
4. Compute the difference between the segmentations (for example, the same DICE loss) and back-propagate through both Siamese sub-networks

The consistency loss is only intended to correct the segmentation and should not overpower the existing labels. Therefore, we begin by training with the segmentation loss only and once the network is sufficiently advanced, we add the consistency branch and perform 2 epochs of segmentation training for each epoch of consistency training. In our experiments, alternating training was essential for good performance: catastrophic forgetting occurred when the network was first trained with segmentation loss and then fine-tuned with consistency loss only.

Note that our approach is not limited to the U-net as a backbone. We use it as it is very popular and easy to train, but generally speaking, any segmentation network can be extended in this fashion.

## 3. DATA

We demonstrate our method on two very different datasets. The first is label-free and has been obtained by holotomographic imaging of live cells on a 3D Cell Explorer-fluo microscope (Nanolive). We use the HeLa ’Kyoto’ cell line [17]. For quantitative evaluation a human expert labeled nucleoli in 15 movie frames (see Fig.2(a) for a snapshot of expert annotations). For the acquisition of training data we used cells with fluorescently tagged nucleoli or histone proteins and imaged them at single time points, with the marker proteins for the different structures imaged in parallel as a single z-slice using FITC or TRITC filters. This z-slice was segmented by the Pixel Classification workflow in ilastik [18] to obtain nucleoli segmentation groundtruth. For nuclei groundtruth we used a pre-trained StarDist network provided by the authors and merged the segmented instances into a foreground/background mask. 98 still frames with corresponding fluorescent markers were used for the nucleoli segmentation task, while 83 were used for the nuclei. Additionally, we acquired two label-free movies with 85 and 260 frames, showing both nuclei and nucleoli.

**Fig. 2.**
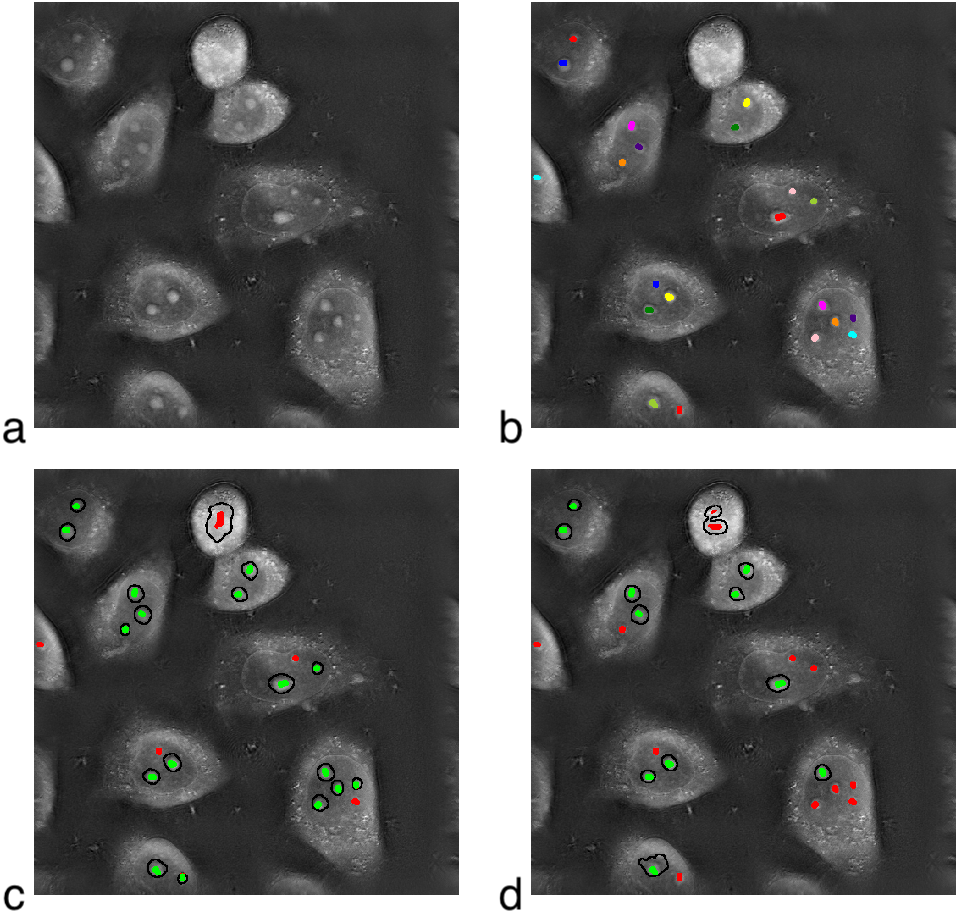
Nucleoli segmentation. (a) Raw data, single video frame, (b) expert annotations as a dot in nucleoli centers, (c) consistent model predictions, (d) baseline predictions. Correct predictions are shown in green, false negative and false positive errors in red.

Improvement of the label-free live imaging segmentation has been the main objective in the development of our method. Nevertheless, as the resulting approach turned out not to be specific for this use case, we performed a second set of experiments on public data from the 2018 Data Science Bowl ([6], still frames) and the Cell Segmentation Benchmark ([8], Fluo-N2DL-HeLa videos).

## 4. EXPERIMENTS AND RESULTS

A holotomographic image consists of 96 2D planes, the majority of which are out of focus. For training data, the in-focus plane is known as it corresponds to the acquired fluorescent groundtruth. For validation and test, we train a simple CNN to find the in-focus plane based on 40 image stacks where the in-focus plane was labeled manually. The CNN consists of 4 residual blocks with a sigmoid activation at the end, trained to predict the probability of each frame to be in focus with a binary cross entropy loss.

A sub-stack of 7 planes above and below the in-focus plane was cropped out of each holotomographic stack to serve as input to the Siamese network (see also Fig. 1). The training was performed alternating 7 batches with segmentation loss and 5 batches with temporal consistency loss. ADAM optimizer was used, the training was stopped after 500 epochs. The resulting foreground/background probability maps were binarized by Otsu thresholding, individual objects were extracted by applying opening and connected components filtering.

First, we verified that the segmentation results did not deteriorate compared to the baseline of training with the segmentation loss only. As the expert labels are object detections rather than segmentations, we evaluate the performance at the object level, computing Average Precision for both networks (exactly as in [7]). On independent frames the proposed Siamese network performs slightly better than the baseline (0.720 vs 0.701 average prediction score). An example evaluation on one of the expert-labeled video frames is shown in Fig. 2. Overall, for video data the Siamese network gives much better predictions than the baseline (0.628 vs 0.512). If the annotated part of the video is not excluded from the unsupervised part of the training – a likely setting for a real-world application of our approach – the average precision score reaches 0.68. The same effect can be observed for the task of nuclei segmentation. For this task we do not have expert labels, so we evaluate the consistency directly. Fig. 3(top) plots the area change for the foreground class in one of the videos. The consistent model exhibits a lot less abrupt changes than the baseline, where the nuclei often ”leak out” to the surrounding cytoplasm for a few frames (Fig. 3(b)).

**Fig. 3.**
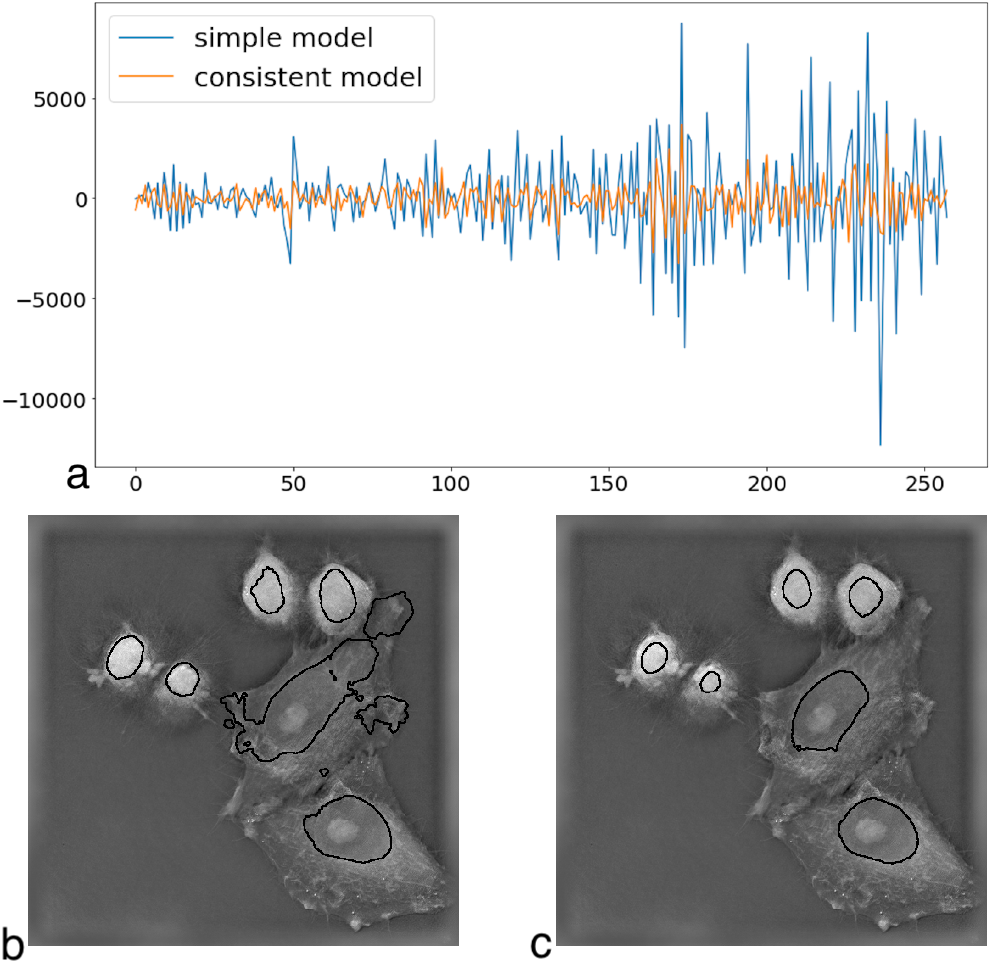
Nuclei segmentation. (a) Change of the predicted total foreground area across time for a movie segmented with both networks (slow biological process, no substantial change expected), (b) a characteristic error of the baseline model, (c) consistent model segmentation of the same frame.

Our 2nd set of experiments aims to show that the applicability of our approach is not limited to holotomographic imaging, as the same method can be applied to improve performance of any fully convolutional segmentation network. To that end, we modify the popular StarDist algorithm [7] in the same manner, making it learn - in addition to segmentation of 2D images - to take into account temporal consistency when segmenting unlabeled videos. We use the same annotated images from the Data Science Bowl as in [7] and fluorescent videos from the Cell Tracking Challenge (Fluo-N2DL-HeLa). The segmentation annotations in the CTC datasets are only used for algorithm evaluation. The temporal consistency loss is realized as the DICE loss on the object center probability channel. We experimented with extending this loss to other channels which predict the direction to object boundary, but this extension did not improve the results compared to using only the object center prediction. Fig. 4 shows qualitative improvement brought by the temporal consistency loss compared to ”standard” StarDist. We observe the same behavior of reduced object ”flickering” in the consistent model as we did for the holotomographic imaging experiments. Fig. 5(a) shows the improvement quantitatively, while Fig. 5(b) confirms that the performance on the original annotated dataset does not deteriorate.

**Fig. 4.**
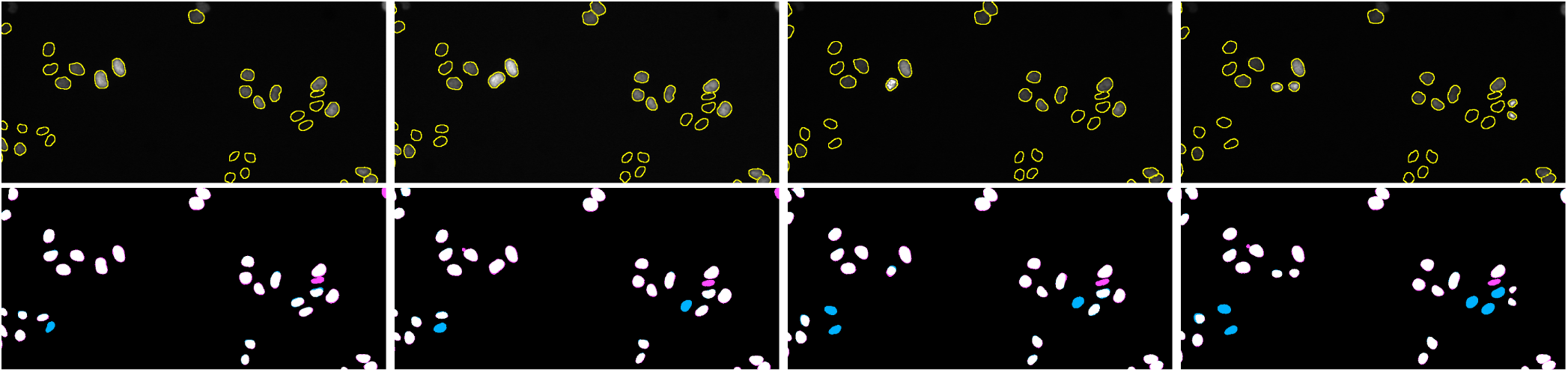
Top: 4 consecutive frames of CTC Fluo-N2DL-HeLa dataset with groundtruth as yellow contours. Bottom: white objects found by both networks, cyan objects missed by the baseline StarDist, magenta objects missed by the consistent StarDist model.

**Fig. 5.**
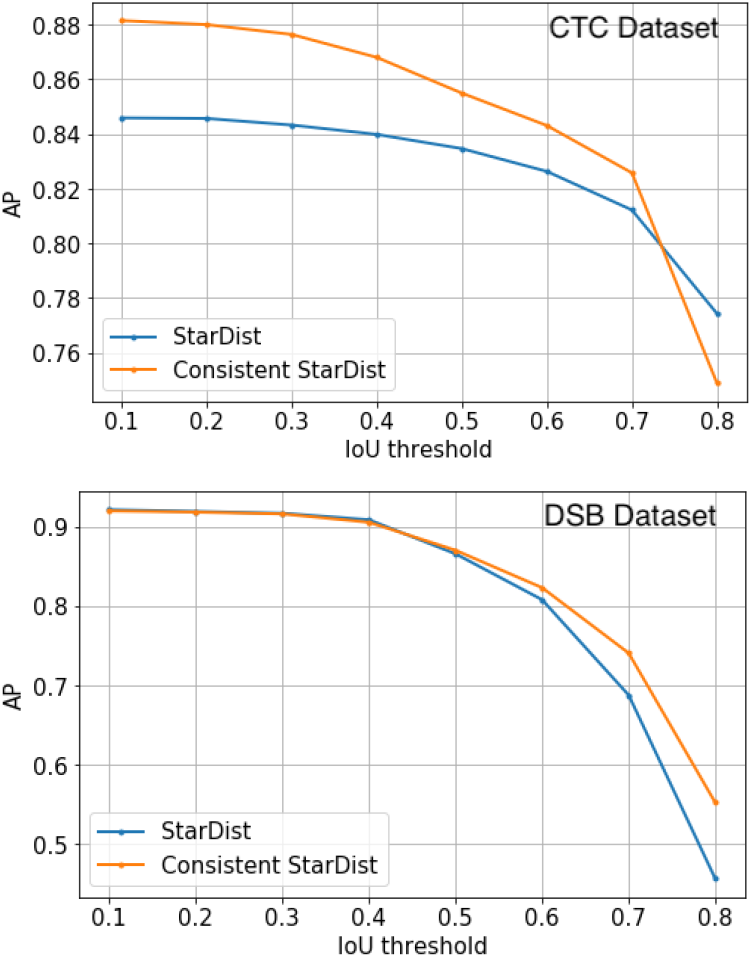
Average Precision score of consistent StarDist network on the DSB (individual images) and CTC (videos) datasets. The network performance on CTC improves with no ill effects on the performance for DSB.

## 5. DISCUSSION

We introduced an approach which allows to improve a semantic segmentation CNN for prediction on videos without annotations. Temporal consistency is induced by an additional loss which penalizes dissimilarity between segmentations of consecutive frames. Segmentation and consistency losses are combined through a Siamese duplication of the CNN. We used DICE loss in our experiments, but other measures of segmentation similarity can be used as drop-in replacement. For example, while DICE would not be applicable to fast moving objects, we can use a loss on the total number of foreground pixels to maintain consistency in this case. Finally, the same approach can be extended to other tasks where groundtruth is only available as independent 2D images, such as segmentation of unlabeled 3D volumes.

## 6. COMPLIANCE WITH ETHICAL STANDARDS

This research study was conducted on publicly available data [6, 8] or generated from previously described cell lines [17]. No ethical approval was required.

## 7. ACKNOWLEDGMENTS

No funding was received for conducting this study. The authors have no relevant financial or non-financial interests to disclose.

